# Social Information Quality and Environmental Volatility Shape Collective Foraging Behavior

**DOI:** 10.1101/2025.11.14.688412

**Authors:** Valerii Chirkov, Ralf H.J.M. Kurvers, Dominik Deffner, Pawel Romanczuk

## Abstract

Collective foraging is widespread across the animal kingdom, allowing animals to more effectively discover resources. However, collective foragers need to balance a key trade off between private exploration and using social information. Social information can come in very distinct forms, ranging from simple positional cues to complex payoff information. However, how the types of available social cues and environmental volatility shape collective foraging behavior is not well understood. We address this using a spatially-explicit model in which agents track a mobile resource via multi-agent reinforcement learning. Agents choose between random exploration, private tracking, and social attraction. We systematically varied resource volatility and the type of available social cues to analyze their effect on individual and collective behavior. Our results show that the quality of social information dictates the emerging collective behavior. Low-quality social cues (e.g., positions, actions) result in a fragile strategy that is effective in stable environments but fails as volatility increases. Conversely, high-quality social information (e.g., payoffs) enables behavioral diversity: Agents selectively copy others and flexibly change between individual tracking or exploration depending on the environmental volatility. Our findings identify the interplay between information quality and ecological context as an important mechanism governing the emergence of distinct forms of collective behavior from individual decision rules.

## 1 Introduction

A fundamental challenge in collective decision-making is determining when to rely on independent exploration versus when to copy others. Animal social foraging serves as a prime testbed for understanding this dynamic, as individuals must constantly balance the costs of discovering resources through private exploration with the benefits of using socially acquired information from their peers [1–3]. This trade-off between private exploration and the use of social information is a central challenge in behavioral ecology, as relying on others can provide a valuable shortcut but also risks the propagation of outdated or erroneous information [4, 5].

To explain how animals solve this problem, classic models often rely on a fixed set of heuristics like “copy the successful” or “copy the majority” [6–8]. While there is robust evidence for the use of such strategies across many contexts, these traditional models necessarily make specific assumptions about what information social cues provide and specify predefined functional forms for their integration. As a result, they fail to capture the processes of how animals flexibly integrate the rich, multidimensional social information available in their surroundings. Social information can, however, vary from discrete location cues (e.g., observing the location of others) all the way to graded public information (e.g., observing foraging success of others) [9, 10]. For instance, a foraging bird might observe a neighbor’s **presence** (positional cue), see it actively **pecking** (action cue), or witness it successfully **extracting** grubs (payoff-based public information) [11]. The adaptive value of each cue is highly context-dependent. In a stable environment, simple cues may already provide useful social information. However, in volatile environments where resources are ephemeral—such as birds tracking thermal updrafts [12] or predators hunting mobile prey [13]—simple positional cues often become rapidly outdated, losing their validity as reliable indicators of resource location. In these contexts, only more complex social information, such as high-fidelity payoff signals, remains adaptive [14]. A significant gap remains in understanding how individuals learn to weigh these distinct cue types based on their reliability and their ecological context.

Deep multi-agent reinforcement learning (MARL) offers a powerful framework to address this gap. This approach lets the model learn the functional relationships between cues directly from experience, enabling the discovery of novel strategies that researchers might not have anticipated. Treating agents as “smart particles” whose decisions and movements are optimized together [15] allows MARL to generate flexible, adaptive policies through reward maximization. This supports recent efforts to adopt more holistic views on social learning and collective intelligence [16, 17]. Prior work has shown that MARL agents can learn biologically plausible behaviors, such as using social information adaptively [18], converging on collective foraging patterns [19, 20], and spontaneously acquiring social learning heuristics [21]. Our work builds on this by providing a principled simulation-based experimental paradigm to investigate how the quality of social information interacts with environmental dynamics to shape emergent collective foraging behavior.

We model a social foraging scenario where agents track a mobile resource performing a correlated random walk (Fig. 1A), mimicking the directional persistence and path tortu-osity observed in many biological prey species [e.g., 22]. The agents’ behavioral repertoire (Fig. 1B) is inspired by fundamental strategies observed across diverse animal species. Agents can engage in a random walk (**exploration**), a widely observed search pattern when animals lack reliable directional cues [23]. Alternatively, agents can use their private information to **track** the resource. This action is modeled as inherently costly, resulting in a slower effective tracking speed than other actions. This cost operationalizes the energetic and cognitive demands of actively sensing and pursuing a resource, such as a predator stalking prey [24], and reflects behavioral constraints like pausing to sense or orient, a constraint central to gradient-following strategies like infotaxis [25, 26]. Finally, agents can move toward a peer (**social attraction**), a core mechanism of social learning where individuals join others to exploit a resource discovered by other agents [11, 27]. Although we acknowledge that acquiring social information—particularly complex payoff signals—can involve significant cognitive and opportunity costs in nature [1], we here choose to model social acquisition as a relatively low-cost alternative to private tracking. This simplification serves to more clearly isolate how different cue types shape the interplay between individual sensing, social information use, and random exploration (the latter acting as a “zero-information” search baseline), while recognizing that real-world scenarios may involve more balanced or context-specific costs for social learning. This action set allows us to probe the fundamental trade-off between relying on costly individual sensing, using socially acquired information, and engaging in uninformed search. To investigate the optimal group-level strategies that emerge from these trade-offs, our primary MARL approach uses centralized training, in which agents share neural network weights. This method is known to effectively find such group-level solutions (which emerge from optimized individual decision rules). We also explored a fully decentralized version, where each agent learns independently using its own distinct network. We systematically vary both the quality of social cues (from neighbor distance to noisy payoff signals) and the environmental volatility (resource speed), to study how their interplay drives emergent collective foraging behavior.

**Figure 1:**
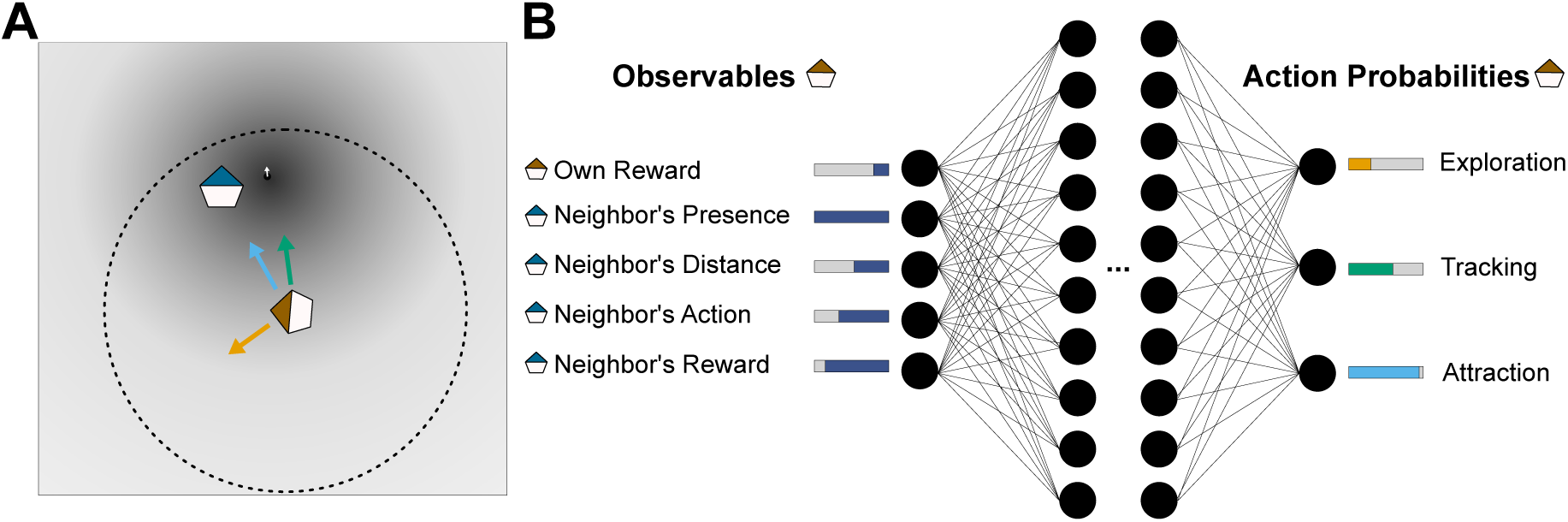
Multi-agent social foraging task. **A**: Schematic of the social foraging scenario. Agents aim to localize and track a continuously moving resource in a 2D environment. The resource consists of a quality gradient (dark-shaded circle with white arrow indicating center and direction of movement; the resource follows a correlated random walk). At each time step, the focal agent (brown) selects one of three discrete actions: random exploration (orange arrow), slow tracking towards the resource’s center (green arrow), or social attraction towards another agent (blue arrow). The focal agent can observe one randomly selected peer (blue agent) within its visual range (dashed circle), and receives rewards based on its proximity to the resource’s center (indicated by the background gradient). **B**: Schematic of the actor network for a single agent. The network maps local, partial observations to action probabilities. Observations include the agent’s own reward, a binary indicator of whether a neighbor is visible, the distance to the observed neighbor, the neighbor’s current action, and the neighbor’s reward. Available social features depend on the current social information condition (see Methods for details).

Our results show that the quality of available social information shapes the emergence of individual-level decision rules as well as the collective performance. And that environmental volatility shapes the adaptive value of social cues. In stable, slow-changing environments, agents limited to low-quality social cues converged on a **Cohesive Tracking** strategy. This strategy achieved near-maximal performance (Fig. 2) by predominantly relying on costly private tracking supplemented with occasional social attraction to maintain a compact cluster (Figs. 3 and 4). However, the performance of this strategy deteriorated quickly when environmental volatility increased as simple social cues became unreliable. Conversely, high-quality payoff information enabled a more robust behavioral repertoire: When private tracking was viable, agents adopted a **Track-or-Copy** strategy, using payoff cues to selectively follow more successful peers – a form of payoff selectivity known to be critical for collective success [7, 14, 28]. As volatility increased and private tracking became ineffective, the group seamlessly transitioned to an **Explore-or-Copy** strategy. This strategy corresponds to a form of distributed collective sensing [12, 29, 30], where agents abandon costly tracking to explore randomly, using payoff cues to aggregate at temporary information hubs created by successful peers (Figs. 3 and 4). Taken together, our results demonstrate that the interplay between information quality and ecological context plays a key role in how individual decision rules give rise to distinct, adaptive forms of collective behavior, generating testable predictions for behavioral research.

**Figure 2:**
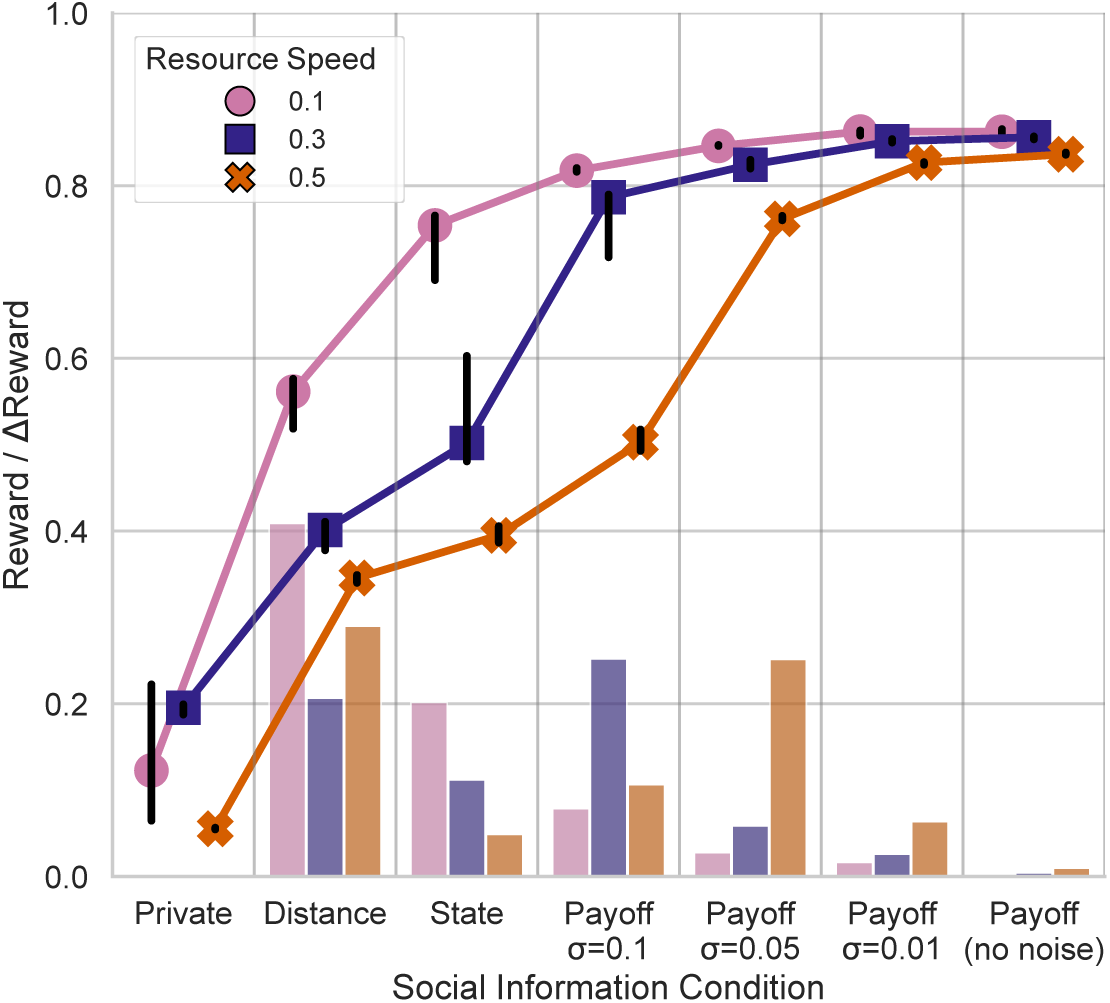
Effect of social information quality on agents’ performance across resource speeds. Median normalized rewards (±95% CI) are shown for agents operating under seven social information conditions, across three resource speeds. Point markers indicate median reward, while bars represent the relative improvement in reward compared to the immediately preceding social information condition. Social information conditions **incrementally** add features: starting from private information only, followed by access to (1) distance to a neighbor, (2) neighbor’s current action (state), and (3) neighbor’s reward signal with decreasing noise (*σ* = 0.1, *σ* = 0.05, *σ* = 0.01, and no noise). Marker shapes and bar colors indicate resource speeds (relative to the constant *v*_max_).

**Figure 3:**
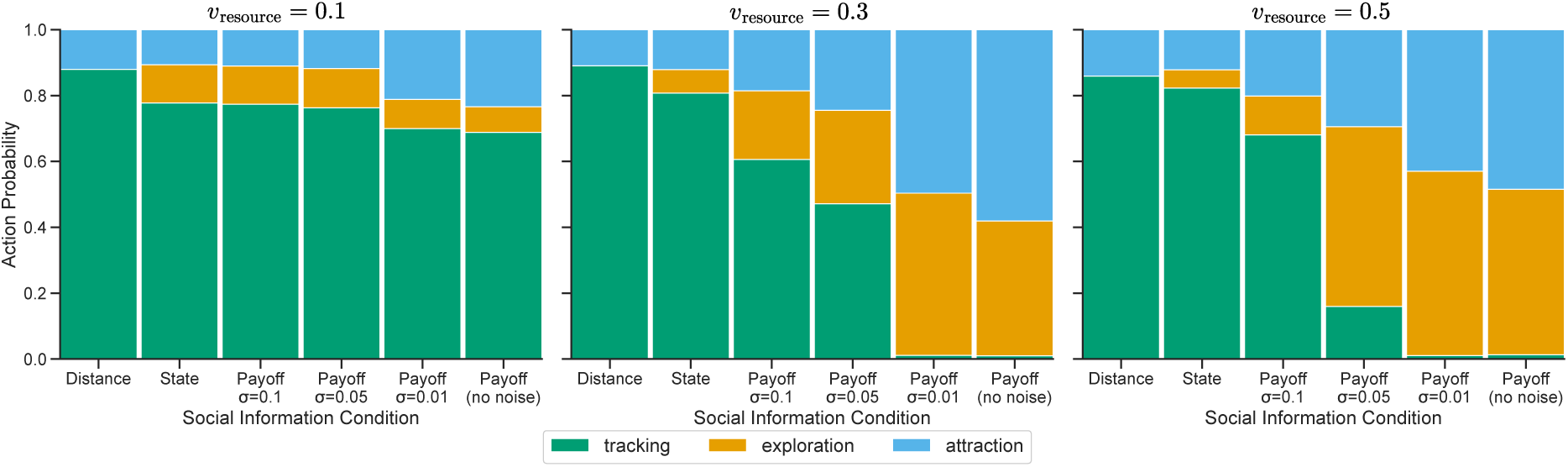
Effect of social information conditions and resource speeds on behavioral repertoire. Bars show the average proportion of time agents spent in each behavioral mode—tracking (green), exploration (orange), and social attraction (blue)—under different combinations of social information and resource speed. Social conditions incrementally add features: starting from private information only, followed by access to (1) distance to a neighbor, (2) neigh-bor’s current action (state), and (3) neighbor’s reward signal with decreasing noise (*σ* = 0.1, *σ* = 0.05, *σ* = 0.01, and no noise). Resource speeds are relative to the constant *v*_max_.

**Figure 4:**
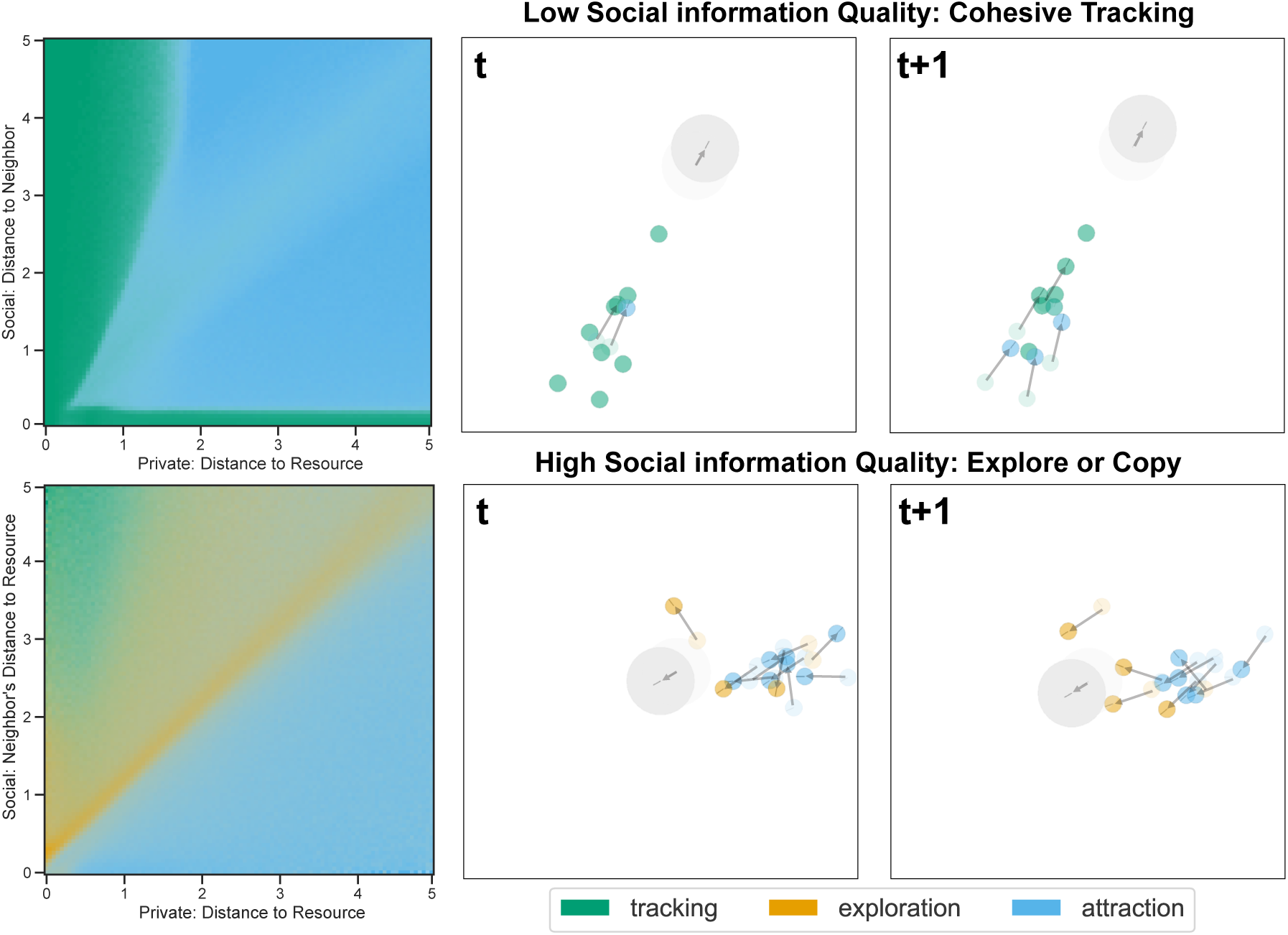
Emergent behavioral rules and group-level patterns. The emergent strategies in the most volatile environment (*v*_resource_ = 0.5 × *v*_max_). The **top row** corresponds to the low-quality “distance-only” condition, showing the Cohesive Tracking strategy. The **bottom row** corresponds to the high-quality “noiseless payoff” condition, showing the Explore-or-Copy strategy. **Left column**: Action probability maps showing the median empirical action frequencies recorded during evaluation rollouts. Data were discretized into 2D bins by observation state, with the median frequency calculated across training seeds and evaluation points, plotted as a function of private distance to the resource’s center (x-axis) and social feature (y-axis). Color intensity reflects the probability of each action: tracking (green), exploration (orange), and social attraction (blue). Each action is plotted as a separate color layer, and the opacity of each cell corresponds directly to the median probability of that action. **Middle and right columns**: Successive frames (*t* and *t*+1). Agent colors mark current behavioral states; vectors from their centers show velocity. The large gray disk indicates the moving resource (its size is for illustration only, as the actual resource provides a gradient covering the entire environment). The *t*−1 and *t* frames (semitransparent) are superimposed on the *t* and *t*+1 frames, respectively, to highlight motion (arrows). Frames are cropped to the active region of the environment. See Supplemental Materials for full videos.

## 2 Materials and Methods

### 2.1 The Collective Foraging Model

#### 2.1.1 Environment and Resource Dynamics

To investigate how social information quality and environmental volatility shape the emergence of individual-level decision rules, we frame collective foraging as a *resource-tracking* problem where agents are rewarded for tracking a mobile resource (Fig. 1A). The task takes place in a continuous 20 × 20 unit square arena *A* with reflective boundaries. A group of *N* = 10 identical agents tracks a single mobile resource for *T* = 1000 discrete time steps.

Let **x***_R_*(*t*) ∈ *A* be the resource’s position at time *t*. The movement of the resource follows a correlated random walk. Turning angles *θ* are sampled uniformly from [0, 2*π*). Step lengths *ℓ* are drawn from a truncated Pareto distribution with a shape parameter *α* = 1 and 1 ≤ *ℓ* ≤ 100. The resource travels at a constant speed *v_resource_* which controls the task’s overall difficulty (see Section 2.3). This truncated power-law distribution generates occasional long relocations, mixing local diffusion with long-distance movements. While not a formal Lévy walk (as truncation ensures finite moments), this pattern characterizes real-world animal and resource movements across a wide range of taxa [31–33]. Tracking this erratic target encapsulates the ecological challenge of following spatiotemporally dynamic resources that underlies various types of foraging [24, 34].

All agents and the resource were initialized randomly for each training and evaluation episode within a 10 × 10 area in the center of the environment to prevent agents and the resource from starting near the arena boundaries.

#### 2.1.2 Agent Dynamics and Reward

Let **x***_i_*(*t*) ∈ *A* be the position of agent *i* at time *t*. Agents can move at a maximum speed of *v*_max_ = 0.05 units per step. The absolute value of this speed is less critical than its value relative to the resource speed (Section 2.3.1) and the effective tracking speed (Section 2.2.1). At each time step, agent *i* receives an individual reward *r_i_*(*t*) based on its proximity to the resource:

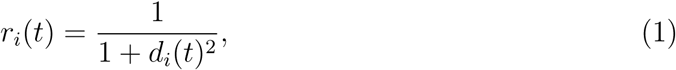

where *d_i_*(*t*) = ‖**x***_i_*(*t*) − **x***_R_*(*t*)‖ _2_ is the Euclidean distance to the resource. This function provides a strong, graded incentive for agents to minimize their distance to the resource.

### 2.2 Agent Behavioral Model

#### 2.2.1 Agent Actions

To probe the fundamental trade-off between private exploration and social exploitation, agents can, at each time step, choose one of three actions, *a_i_*(*t*) ∈ {Exploration, Tracking, Social Attract (Fig. 1B). The chosen action dictates the agent’s movement direction, i.e. the direction of its current velocity vector **v***_i_*(*t*). The position is updated in discrete time by **x***_i_*(*t* + Δ*t*) = **x***_i_*(*t*) + **v***_i_*(*t*)Δ*t* with Δ*t* = 1, subject to reflection at arena boundaries. The actions represent distinct foraging strategies observed in natural systems:

1. **Exploration** models random search behavior when reliable directional cues are lacking. The agent performs a correlated random walk at maximum speed (*v*_max_). The movement direction *ϕ_t_* is updated by *ϕ_t_* = *ϕ_t−_*_1_ + Δ*ϕ*, where the turning angle Δ*ϕ* is sampled from a von Mises distribution (a circular analogue of the Gaussian distribution commonly used to model animal orientation; [see e.g. 35, 36]), Δ*ϕ* ∼ vonMises(*µ* = 0*, κ* = 2). The resulting movement vector is **v***_i_*(*t*) = *v*_max_ · [cos *ϕ_t_,* sin *ϕ_t_*].
2. **Tracking** represents costly exploitation based on private information. The agent attempts to move towards the center of the resource. Let **u**_track_(*t*) = (**x***_R_*(*t*) − **x***_i_*(*t*))*/d_i_*(*t*) be the unit vector toward the resource. The movement is stochastic to operationalize the costs associated with acquiring and acting on private information (e.g., pausing to sense, orient, and handle prey [25]):

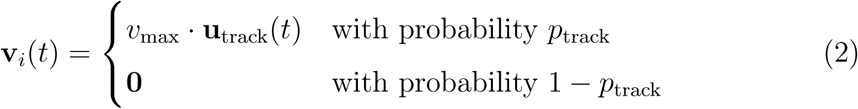 The probability 1 − *p*_track_ of remaining stationary can be interpreted as the temporal cost of selecting the tracking action. A higher cost (larger 1 − *p*_track_) reduces the agent’s effective movement speed when tracking, yielding an effective average tracking speed of *v*_tracking_ = *p*_track_ × *v*_max_. For our main experiments, *p*_track_ = 0.1, resulting in *v*_tracking_ = 0.1 × *v*_max_. The cost parameter 1 − *p*_track_ was varied in robustness analyses (see Section 2.5).
3. **Social attraction** represents leveraging social information by moving towards peers, potentially avoiding the cost of slow tracking, a core mechanism of social foraging [1]. The agent moves at maximum speed (*v*_max_) toward its currently observed peer *j*. Let **u**_social_(*t*) = (**x***_j_*(*t*) − **x***_i_*(*t*))*/*‖**x***_j_*(*t*) − **x***_i_*(*t*)‖_2_ be the unit vector toward the peer. The movement vector is **v***_i_*(*t*) = *v*_max_ · **u**_social_(*t*).

The “Tracking” and “Exploration” actions were always available. “Social Attraction” was only meaningful if a peer was within the visual range. To maintain a consistent action space while preventing the policy network from processing undefined inputs when social cues were absent, we introduced a penalty mechanism. If an agent selected “Social Attraction” when no peer was visible (due to visual range limits or the experimental condition), it received a large penalty *r*_penalty_ = −1, and its movement direction remained unchanged (**v***_i_*(*t*) = **v***_i_*(*t*−1)). This ensures the policy learns to avoid inapplicable actions. Such invalid selections are rare, typically occurring only early in training or in edge cases with limited visibility, and this penalty mechanism had negligible impact on the final learned strategies.

#### 2.2.2 Agent Observations

Agents operate under partial observability, receiving only local information. To model both private information and the different types of social cues (distance, action, payoff), each agent *i* perceives its environment through a five-dimensional observation vector **o***_i_*(*t*), showed schematically in Fig. 1B. This is constructed based on a visual range *r*_vis_ = 15 units (see Section 2.5 for robustness checks).

Let *V_i_*(*t*) = {*j ≠ i* | ‖**x***_j_*(*t*) − **x***_i_*(*t*)‖_2_ ≤ *r*_vis_} be the set of peers visible to agent *i* at time *t*. To model perceptual and attentional constraints often observed in animal groups [e.g. 37], and to simplify the analysis by avoiding complex social information aggregation rules, if *V_i_*(*t*) is not empty, agent *i* attends to *exactly one* peer *j*, sampled uniformly at random from *V_i_*(*t*) at each time step. While animals in nature typically bias their attention toward closer neighbors, this uniform sampling—bounded by a maximum visual range—simplifies our analysis by avoiding complex distance-weighted aggregation rules. The agent then uses this focal peer’s information to decide whether to approach it (via the Social Attraction action) or choose a different behavior. If *V_i_*(*t*) is empty, no social information is gathered. Because the focal neighbor is resampled independently at each time step, the attention process is ergodic: given enough time, an agent will observe each group member with equal frequency. This corresponds to an effective averaging of interactions across neighbors over time. Thus, our qualitative results are representative of a setting where each agent observes *all* neighbors simultaneously in the limit of a slowly changing environment, analogous to results on synchronization in blinking network models [38, 39].

The observation vector **o***_i_*(*t*) = (*o*_1_*, o*_2_*, o*_3_*, o*_4_*, o*_5_) components represent different information types:

1. *o*_1_ = *r_i_*(*t*): Private information about own success (i.e., reward).
2. *o*_2_ = I(*V_i_*(*t*) ≠ ∅): Basic social cue indicating neighbor presence.
3. *o*_3_ = *d_ij_*(*t*): Positional social cue (distance to observed peer *j*). Set to 0 if *o*_2_ = 0.
4. *o*_4_ = *a_j_*(*t* − 1): Behavioral social cue (observed peer *j*’s previous action). Set to 0 if *o*_2_ = 0.
5. *o*_5_ = *r̃_j_*(*t*): Payoff-based public information (observed peer *j*’s previous reward), corrupted by additive Gaussian noise *ϵ* ∼ N (0*, σ*^2^) to model imperfect observation. *r̃_j_*(*t*) = *r_j_*(*t*) + *ɛ*. Set to 0 if *o*_2_ = 0.

### 2.3 Simulation Design

To address the central question of how resource volatility interacts with the quality of social information, we systematically varied two factors: Resource Speed (proxy for environmental volatility) and Social Information Quality (controlling the type and fidelity of social cues).

#### 2.3.1 Resource Speed

To simulate environments ranging from stable to volatile, we varied resource speed (*v*_resource_) relative to the agents’ maximum speed (*v*_max_) across three levels:

- **Slow:** Resource speed 0.1 (*v*_resource_ = 0.1 × *v*_max_).
- **Medium:** Resource speed 0.3 (*v*_resource_ = 0.3 × *v*_max_).
- **Fast:** Resource speed 0.5 (*v*_resource_ = 0.5 × *v*_max_).

#### 2.3.2 Social Information Quality

To explore the value of different social cues (distance, action, payoff) and the fidelity of the payoff signal, we designed seven experimental conditions that progressively increased the information available in the agent’s observation vector (**o***_i_*(*t*)). Starting with purely private information and progressively adding more complex social cues:

1. **Private:** Only *o*_1_ (own reward) available.
2. **+Distance:** *o*_1_*, o*_2_*, o*_3_ available (reward, presence, distance).
3. **+Action:** *o*_1_*, o*_2_*, o*_3_*, o*_4_ available (reward, presence, distance, peer action).
4. **+Payoff (High Noise):** All components *o*_1_*, … , o*_5_ available, with high noise on peer reward *o*_5_ (*σ* = 0.1).
5. **+Payoff (Medium Noise):** All components available, with medium noise on *o*_5_ (*σ* = 0.05).
6. **+Payoff (Low Noise):** All components available, with low noise on *o*_5_ (*σ* = 0.01).
7. **+Payoff (No Noise):** All components available, with no noise on *o*_5_.

### 2.4 Multi-Agent Reinforcement Learning (MARL)

#### 2.4.1 Training Algorithm

We adopt a MARL approach to train agents to maximize their individual rewards, allowing collective strategies to emerge. Specifically, we used Multi-Agent Proximal Policy Optimization (MAPPO) [40, 41], a widely used algorithm based on the centralized training–decentralized execution (CTDE) paradigm. CTDE is well-suited for partially observable multi-agent problems; the centralized critic leverages global information during training to stabilize learning and improve coordination [42], while the decentralized actors learn policies based only on local observations, ensuring scalability during execution. All agents shared the parameters of a single actor policy network and a single centralized critic network.

#### 2.4.2 Policy and Critic Architecture

Both the actor (policy) and critic (value) networks were multi-layer perceptrons (MLPs) with two hidden layers of 256 units each, using tanh activation functions. The actor network mapped local observations **o***_i_*(*t*) to probabilities for the three actions via a final sigmoid layer and a categorical distribution (see schematic illustration in Fig. 1B). The critic network mapped the joint state (approximated using all agents’ local observations during centralized training) to a scalar state-value estimate. We utilized standard PPO components: a clipped surrogate objective for the policy loss, a mean squared error loss for the value function, and an entropy bonus to encourage exploration. Advantages were calculated using Generalized Advantage Estimation (GAE) [43] applying advantage normalization and value clipping. Key training hyperparameters are detailed in Supplementary Information Table S2.

#### 2.4.3 Training and Implementation Details

The simulation environment was implemented using the Vectorized Multi-Agent Simulator VMAS v1.4.3 [44]. All MARL training was conducted using PyTorch v2.5.1 [45] and the TorchRL library v0.6.0 [42], using its TensorDict v0.6.2 data structures. A separate policy was trained from scratch for each unique combination of Resource Speed, Social Information Quality, and training seed. Each training run encompassed 480 iterations, gathering 500000 environment frames per iteration (equivalent to 500 episodes of 1000 steps). This resulted in a total of 240 million environment interactions per run. Training was performed on a single NVIDIA A100 or V100S GPU, with an average runtime of 17.9 hours (std=1.8). All runs used distinct random seeds for initialization and data generation. Training metrics were logged using wandb v0.19.2 [46]. Figures were generated using Matplotlib v3.9.0 [47] and Seaborn v0.13.2 [48]. For details of the third-party software see Supplementary Information Table S1.

### 2.5 Robustness and Sensitivity Analyses

To assess the robustness of the emergent strategies and the sensitivity of our findings to specific modeling choices, we conducted several additional sets of experiments.

- **Tracking Cost / Effective Speed:** We systematically varied the tracking cost 1 − *p*_track_ (Eq. (2)) to explore how the intrinsic cost of private information affects strategy emergence. We tested three levels: 1 − *p*_track_ ∈ {0.7, 0.8, 0.95}, corresponding to effective tracking speeds *v*_tracking_ ∈ {0.3 × *v*_max_, 0.2 × *v*_max_, 0.05 × *v*_max_}.
- **Decentralized Training:** To ensure our results were not dependent on the centralized training setup, we replicated the main experiments using a fully-decentralized training algorithm (Independent PPO, IPPO [49]). In IPPO, each agent learns independently using its own actor and critic network, with no parameter sharing or access to other agents’ information during training.
- **Visual Range:** To test sensitivity to perceptual limitations, we repeated the main experiments with reduced visual ranges, setting *r*_vis_ ∈ {10, 5, 1} units, compared to the default *r*_vis_ = 15.

For the Tracking Cost and Decentralized Training analyses, the training procedure, number of seeds (seven), evaluation methodology, and strategy consistency analysis followed the same protocols as for the main experiments. The Visual Range analysis followed the same general training and evaluation procedure but was not reproduced across multiple seeds.

### 2.6 Data Collection and Evaluation Procedure

#### 2.6.1 Performance Evaluation

Following best practices for RL reproducibility [50], each experimental condition was trained using seven independent random seeds. Learned policies were evaluated every 20 training iterations during the final phase of training (iterations 400 to 480) to assess performance near convergence. Each evaluation consisted of 1000 independent rollout episodes per seed. This resulted in 5 evaluation points × 1000 episodes/point = 5000 evaluation episodes per seed. Final performance metrics (e.g., average reward) reported in the Results are averaged across these 5000 episodes and then across the 7 seeds. Error bars represent 95% confidence intervals computed across the 7 independent seeds.

#### 2.6.2 Strategy Consistency Analysis

To confirm that the training process consistently converged to similar behavioral strategies across different random seeds within the same condition, we performed a post-hoc analysis of behavioral strategy similarity. For each condition, we first calculated the average action composition vector (proportion of time spent in Exploration, Tracking, Social Attraction, analogous to the behavioral analysis described in the Results Section 3.2) for each of the 7 seeds, averaging across all 5000 evaluation episodes for that seed. We then computed the pairwise cosine similarity between these 7 strategy vectors.

Agglomerative hierarchical clustering was performed on the cosine distance matrix (1−similarity) derived from these vectors. We examined the first bifurcation (split at the root) of the clustering tree. If the average cosine similarity *between* the two resulting clusters fell below a threshold of 0.9, we flagged the smaller cluster as representing a divergent or potentially unconverged strategy. In such cases, the runs belonging to the smaller cluster were excluded from the main analysis to ensure reported results reflect the dominant, consistent behavioral mode. This conservative pruning procedure, implemented using SciPy [51] and Scikit-learn [52], affected only 5 out of 147 total training runs (approx. 3.5%), mostly in non-social conditions. Overall consistency within the retained dominant clusters was very high (average cosine similarity *>* 0.95), confirming the robustness of the emergent strategies (for details see Supplementary Information Table S3).

#### 2.6.3 Strategy Visualization

To visualize the agents’ emergent behavioral rules (Figs. 4 and 5), we calculated empirical action frequencies from the evaluation data. Specifically, we recorded the continuous observation states and the corresponding chosen actions across all 1000 evaluation rollout episodes per seed (see Section 2.6.1 for details). The observation space was then discretized into a 2D grid, and the relative frequency of each action was calculated within each bin. The final heatmaps display the median of these empirical frequencies across 5 evaluation points for each independent training seeds. The final visualization (Figs. 4 and 5) is generated via alpha blending, where each action is plotted as a separate color layer. The opacity (alpha channel) of each cell corresponds directly to the median probability of that action.

**Figure 5:**
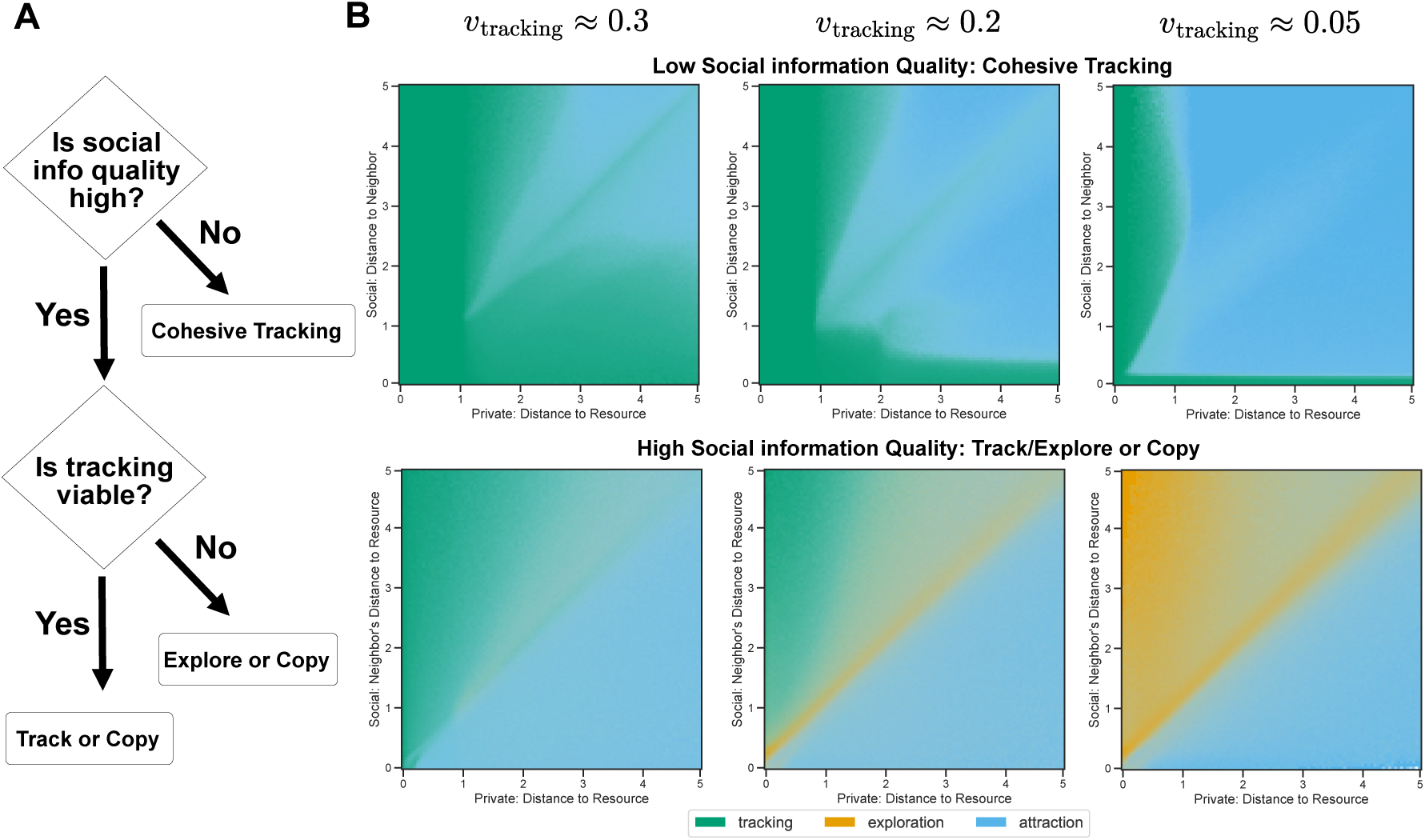
Ecological Conditions Determine the Emergence of Collective Foraging Strategies. **A**: Schematic illustration of how the interplay of social information quality and the viability of private tracking determines which collective strategy emerges. **B**: Action probability maps showing the median empirical action frequencies recorded during evaluation rollouts. Data were discretized into 2D bins by observation state, with the median frequency calculated across training seeds and evaluation points. The maps show the action probabilities for agents in the most volatile environment (*v*_resource_ = 0.5 × *v*_max_) under varying effective tracking speeds (*v*_tracking_ = 0.3 × *v*_max_, 0.2 × *v*_max_, and 0.05 × *v*_max_ from left to right). Colors indicate the probability of tracking (green), exploration (orange), or social attraction (blue). Each action is plotted as a separate color layer, and the opacity of each cell corresponds directly to the median probability of that action. The **top row** (low-quality cues) shows that agents remain locked in the Cohesive Tracking strategy, increasing their use of social attraction as tracking becomes less viable. The **bottom row** (high-quality cues) demonstrates behavioral flexibility, showing a clear transition from a Track-or-Copy strategy (left) to an Explore-or-Copy strategy (right) as tracking viability decreases.

## 3 Results

We present our findings as follows. We first establish that the adaptive value of any given social cue is context-dependent, with high-quality payoff signals becoming essential as environmental volatility increases. We then explore how these performance differences arise from shifts in individual agent behavior, leading to the emergence of collective tracking strategies. Finally, we confirm that these emergent strategies are robust outcomes of the agent-environment interaction by testing their stability against changes in both intrinsic costs and the learning algorithm itself.

### 3.1 The value of social information depends on environmental volatility

To understand how social information quality and environmental volatility shape performance, we first established their combined effect on foraging success. We denote resource speed by *v*_resource_, the normalized reward (i.e., the collected reward divided by the maximum possible reward) by *R*, and changes in the reward by Δ*R*. Unless stated otherwise, we report medians with 95% bootstrap confidence intervals in brackets; full statistics can be found in Supplementary Information Table S4. As predicted, access to higher quality social information consistently improved agent performance, but the magnitude of this benefit was dictated by environmental volatility (Fig. 2).

In the baseline condition with only private information, performance was generally low across all resource speeds. As expected, performance was the lowest in the fast environment. Surprisingly, the median reward for the slow-moving resource (*v*_resource_ = 0.1 × *v*_max_) was lower than for the medium-speed resource (*v*_resource_ = 0.3 × *v*_max_). This was due to the agents in the slower environment converging to different proportions of tracking and exploration across training seeds. This led to larger confidence intervals and a lower overall median reward, even after pruning based on strategy similarity (see Supplementary Information Table S3 for strategy consistency results).

Once social information was introduced, clear and consistent performance differences emerged. In stable environments with a slow-moving resource (*v*_resource_ = 0.1 × *v*_max_), agents gained the most from low-quality social cues. The reward jumped substantially when distance was added (Δ*R* = 0.409[0.367, 0.424]), and again when adding a neighbor’s action state (Δ*R* = 0.202[0.139, 0.214]). Further adding high-quality payoff signals offered only modest improvements, suggesting that in predictable environments, simple social cues are sufficient for effective coordination.

This pattern reversed in volatile environments. For fast-moving resources (*v*_resource_ = 0.5 × *v*_max_), the limitations of low-quality cues became apparent. While neighbor distance still provided an initial benefit (Δ*R* = 0.291[0.285, 0.294]), information about a neighbor’s action was of little value (Δ*R* = 0.049[0.041, 0.060]). Instead, the most substantial performance gains came from access to high-quality payoff signals, with foraging success increasing by Δ*R* = 0.252[0.250, 0.256] when payoff noise was reduced from high (*σ* = 0.1) to moderate (*σ* = 0.05). This highlights that as environmental unpredictability increases, the ability to accurately assess the success of others becomes critical. Interestingly, in all environments, performance saturated before payoff signals became noiseless (minimal gains for *σ <* 0.05), demonstrating that the learned strategies are robust to the moderate levels of informational noise characteristic of natural systems [2, 30].

### 3.2 Adaptive shifts in individual foraging strategy

The performance gains documented above were driven by corresponding shifts in the agents’ individual behavioral strategies. We examined how agents allocated time among the three available actions—tracking, exploration, and social attraction—and found that their collective behavior emerged from context-dependent changes in these individual decisions (Fig. 3). Full statistics can be found in Supplementary Information Table S5.

In stable, slow-moving environments (*v*_resource_ = 0.1), tracking was the dominant behavior across all conditions. However, as environmental volatility increased, the cost-benefit ratio of private tracking versus the other behaviors shifted. This prompted agents to rely more on other behaviors. The specific type of social information available determined how this shift occurred. With access to only neighbor distance (low-quality social cue), agents completely avoided exploration across all three resource speeds (mean exploration = 0[0, 0]; Fig. 3, leftmost bars). We interpret this striking behavioral shift as an emergent solution to the problem of social noise; by collectively ceasing to explore, the group effectively enhances the reliability of the social signal, ensuring that an agent’s position is more strongly correlated with the resource location and thus increasing the value of simple proximity cues [53].

When high-quality payoff signals were available, a more dramatic transformation occurred. Agents sharply reduced their reliance on costly private tracking, instead adopting a mixed strategy of exploration and social attraction. In the most volatile environment (*v*_resource_ = 0.5) with noiseless payoff signals, agents spent almost no time tracking (1.3%[0.010, 0.016]), dividing their efforts almost equally between exploration (50.3%[0.470, 0.536]) and social attraction (48.5%[0.449, 0.518]). This strategic shift enabled a fundamentally different, and more effective, collective response to environmental circumstances.

### 3.3 Emergence of distinct collective behaviors

The shifts in individual behavior give rise to distinct, group-level strategies. To illustrate these findings, Fig. 4 visualizes the learned spatial dynamics and decision rules for these contrasting strategies, while Fig. 5 provides a conceptual summary and demonstrates how strategy selection is driven by tracking viability. As summarized by the conceptual flowchart in Fig. 5A, the quality of social information is the primary determinant of the group’s behavioral flexibility. Low-quality social cues constrain the group to a single strategy, whereas high-quality social cues enable a flexible response based on the viability of private tracking. Below, we detail the three major emergent strategies, focusing on the most volatile environment (*v*_resource_ = 0.5 × *v*_max_).

#### Cohesive Tracking

When agents only have access to low-quality social information (e.g., neighbor distance), they are locked into a single strategy: Cohesive Tracking. In this mode, agents predominantly rely on private tracking while using social attraction non-selectively to maintain group cohesion. The underlying rule is simple: an agent follows a peer unless it is already very close to the resource’s center or the peer itself is very closeby (Fig. 4, top panels; Supplementary Videos 1). This results in a compact, mobile cluster that moves as a coherent unit, reminiscent of collective movements where leaders “pull” followers along [54]. While effective in predictable environments, the strategy’s strong reliance on private tracking means its performance deteriorates when the resource becomes more volatile (Fig. 3) or the costs of tracking increase due to lower effective tracking speed relative to resource speed (Fig. 2 and Fig. S2).

#### Track-or-Copy

When high-quality payoff information is available and private tracking is viable (i.e., tracking cost is low due to greater effective tracking speed relative to resource speed), a more straightforward Track-or-Copy strategy emerges. Here, agents still default to private tracking but now use social information selectively. By observing a neighbor’s payoff at each time step, they follow a rule akin to the “copy the successful” heuristic [7]: track privately, but if a more successful neighbor with a higher payoff is detected, copy them. This decision rule is clearly visible as a diagonal boundary in their action probability map (Fig. 5B, bottom-left panel; Supplementary Videos 2): agents choose social attraction only when the observed peer is closer to the resource center (i.e., has a higher payoff) than themselves; otherwise, they default to private tracking. This represents a more efficient use of social information than simple cohesion.

#### Explore-or-Copy and Distributed Sensing

When high-quality payoff information is available but private tracking is rendered ineffective by high environmental volatility (i.e., the cost of tracking increases due to lower effective tracking speed relative to resource speed), the group flexibly switches to an Explore-or-Copy strategy. While this strategy uses the same social rule as Track-or-Copy (i.e., copy a more successful peer), it is fundamentally different in its default asocial behavior. Agents abandon costly tracking altogether and instead engage in random exploration. Put simply, if a neighbor with a higher payoff is detected, copy them; if not, explore randomly (Fig. 4, bottom panels; Fig. 5B, bottom-right panel Supplementary Videos 3). This individual rule gives rise to a powerful emergent behavior that functions as a form of distributed collective sensing [12, 29, 30]. Successful explorers become temporary information hubs, attracting other agents and allowing the group as a whole to effectively track the resource in a decentralized manner.

### 3.4 Robustness of Emergent Strategies

To further test the robustness of our findings, we conducted three additional sets of experiments where we manipulated key aspects of our model: the intrinsic cost of tracking, the training algorithm itself, and the visual range.

#### Effective Tracking Speed

The main results used a fixed *effective tracking speed v*_tracking_ = 0.1 × *v*_max_ (resulting from *p*_track_ = 0.1). To test the robustness of our findings, we systematically varied the effective tracking speed, testing levels *v*_tracking_ ∈ {0.3 × *v*_max_, 0.2 × *v*_max_, 0.05 × *v*_max_} (corresponding to *p*_track_ ∈ {0.3, 0.2, 0.05}). Not surprisingly, a lower effective tracking speed (i.e., higher tracking cost) disproportionately harms performance in low-quality social information conditions (“distance” and “action”), as agents in these scenarios rely heavily on the Cohesive Tracking strategy (Fig. S2). As the effective tracking speed is reduced (i.e., tracking cost is increased), tracking is progressively “pushed out” of the agents’ learned policies and replaced by exploration and attraction (Fig. 5B). With high-quality social information, this leads to a complete strategic shift: for a high tracking speed (*v*_tracking_ = 0.3 × *v*_max_), Track-and-Copy is preferred, but for a low tracking speed (*v*_tracking_ = 0.05 × *v*_max_), agents switch entirely to Explore-and-Copy (Fig. 5B; Fig. S3). The fact that the behavioral transitions caused by increasing the internal tracking cost (lowering *v*_tracking_ in Fig. 5B) mirror those caused by increasing external environmental volatility (higher *v*_resource_ in Fig. 3) confirms that the fundamental driver of strategy selection is the overall viability of private tracking, whether limited by the environment or by the organism’s own energetic and cognitive constraints.

#### Decentralized Training

The main results used a centralized training procedure (MAPPO) known to foster coordination. To test if our findings depend on this, we reproduced our results using a fully decentralized algorithm (IPPO) where each agent learned fully independently (see Methods section for more information). The same fundamental strategies emerged (Fig. S5B-C), confirming the robustness of the core relationships between social information quality, environmental volatility, and strategy selection. However, decentralized training led to a noticeable drop in performance, particularly in low-quality information conditions (Fig. S4) and an increased reliance on social cues across all settings (Fig. S5A). Interestingly, the learned policies in the ‘distance-only’ condition under decentralized training showed a stronger prioritization of tracking over social attraction compared to the centralized case (compare Fig. 4 top-left and Fig. S5B). This suggests that the Cohesive Tracking strategy, which relies on maintaining a tight cluster, may require a degree of cooperation facilitated by centralized training, while decentralized learning appears to favor individual tracking, potentially at the expense of group cohesion. These findings highlight that Cohesive Tracking is sensitive not only to ecological conditions, but also to learning dynamics.

#### Visual Range

We evaluated the agents’ performance under increasingly restricted perceptual limits. Performance remained largely unchanged when the visual range was reduced to *r*_vis_ = 10 and *r*_vis_ = 5, indicating that the emergent strategies are robust to moderate perceptual constraints. Performance deteriorated only at the extreme minimal range of *r*_vis_ = 1 (full results detailed in Supplementary Information Section S1).

## 4 Discussion

Our study demonstrates how environmental volatility and the availability and type of social information interactively shape the emergence of collective foraging strategies. By systematically varying the type and fidelity of social features in a simulated social foraging task, we showed that agents adaptively integrate social cues based on their predictive value within specific ecological contexts. We identified two distinct behavioral regimes determined by information quality. The first is a fragile, cohesive strategy reliant on low-quality social cues. The second is flexible and adaptive, emerging only when high-quality social information is available.

With only access to low-quality social cues (neighbor proximity or action), agents converged on a Cohesive Tracking strategy. This strategy, effective in stable, slow-moving resource environments, relies on all agents predominantly using costly private tracking while using social cues to maintain a cohesive group. However, this strategy was not robust. Its success was highly sensitive to ecological context; performance degraded significantly as environmental volatility (resource speed) increased or the cost of private tracking increased (i.e., lower effective *v*_tracking_; see Fig. 5B).

In sharp contrast, access to high-quality payoff information allowed the group to be more flexible in their behavior. This enabled them to switch strategies based on the viability of private tracking. When private tracking was effective (slow resource or low tracking cost), agents adopted a Track-or-Copy strategy, using social information to selectively copy more successful peers. When private tracking became ineffective (fast resource or high tracking cost), the group evolved an Explore-or-Copy strategy. This emergent behavior functions as a form of distributed sensing, where agents abandon costly tracking, explore randomly, and use payoff cues to aggregate at temporary information hubs created by successful explorers (Fig. 4).

The emergence of the “Explore-or-Copy” strategy highlights a flexible dynamic related to the classic Producer-Scrounger (P-S) dilemma [1, 55–58]. It is important to note that traditional P-S models typically assume stationary, depleting resources where individuals actively compete, creating a strict dilemma around information sharing. While our model features a mobile, non-depleting resource–thereby removing direct competition over the resource itself–it nonetheless captures the core informational trade-off of the P-S framework. In our model, agents are not locked into fixed “Producer” (Exploration/Tracking) or “Scrounger” (Social Attraction) roles. Rather, the availability of high-fidelity payoff information enables them to dynamically adopt the most profitable role at any given time. An agent can act as a producer by exploring. The instant an agent observes a more successful peer, they switch to the scrounger role to copy that peer. This emergent behavior closely mirrors empirical evidence showing that animals conditionally rely on social information during foraging, by primarily using it when they are unsuccessful themselves [6, 59–62]. Such selectivity makes strategies based on high-quality social information not only robust to ecological factors (like volatility) but also robust to social uncertainty. The group becomes effectively “immune” to the dilemma’s primary pitfall: the risk of scroungers copying other unsuccessful scroungers, leading to negative information cascades [53, 63]. Because successful agents are reliably identified, the incentive to explore (produce) is maintained. In natural systems, however, assessing the performance of others may entail its own time and cognitive costs. Dynamic switching is possible only because high-quality social information enables agents to differentiate successful peers from unsuccessful ones. When payoff information is noisy, this filtering capacity breaks down. This makes agents vulnerable to copying errors, which degrades performance. The highly flexible, Explore-or-Copy strategy contrasts sharply with the Cohesive Tracking strategy, which appeared to require significant group-level coordination to maintain a tight formation, forcing agents to prioritize group cohesion over immediate, individual tracking. Our decentralized training experiments supported this, showing that performance in this low-quality social information regime dropped significantly without a mechanism to establish this group-wide behavioral consensus.

The emergence and viability of these strategies depend on the assumption that individual tracking is more costly than using social information. This asymmetry is essential to generating the central trade-off that we aimed to study. If social learning were equally or more expensive than private tracking, the incentive to copy others would disappear, resulting in purely asocial behavior. It is important to note, however, that operationalizing these relative costs when modeling biological systems is crucial, yet highly non-trivial. “Cost” can be expressed in various ways: as temporal delays (as implemented in our study), reduced precision of private information, or direct energetic costs. At the same time, acquiring high-fidelity public information, such as accurately assessing a conspecific’s exact payoff, could also impose its own significant perceptual and cognitive demands in many naturalistic scenarios. If these social acquisition costs outweigh the energetic burden of private sensing, we would expect a higher threshold for using social information, which would likely drive groups toward independent search strategies.

From a broader theoretical perspective, the learned strategies functionally encode the value of information, reflecting an adaptive valuation based on its utility in reducing uncertainty [2, 64]. In our model, simple, low-bandwidth cues–analogous to observing a neighbor’s mere presence–are sufficient to reduce uncertainty in stable environments; indeed, even when high-bandwidth, high-fidelity signals–analogous to observing a peer’s actual foraging success–are available, private tracking is still preferred. However, as environmental volatility increases, relying on simple cues becomes less adaptive, and high-fidelity signals become necessary to sustain collective performance. It has been argued, however, that in many natural systems, animals often only have access to simple behavioral cues rather than explicit, payoff-based information, making groups vulnerable to suboptimal informational cascades [e.g., 53]. Our results show that when high-quality social cues are unavailable and the environment is highly volatile, social information becomes fundamentally unreliable. In such contexts, the best an individual can do is refrain from using social cues entirely and rely instead on private information. Our findings thus mechanistically demonstrate that adaptive foraging requires constantly balancing information sources–a complex trade-off dictated by the specific types of available social cues and the volatility of the environment.

Using deep reinforcement learning, our study establishes a normative baseline for navigating these complex trade-offs optimally. While a single animal within its lifetime would not experience the millions of interactions necessary for our agents to learn artificial network weights, our goal was not to simulate precise biological ontogeny. Rather, our framework acts as an optimization tool to reveal which collective strategies maximize fitness under specific constraints. In nature, these highly adapted solutions could be approximated through a combination of evolutionary adaptation and heuristic-based learning.

We investigated optimal social information usage in a simplified environment with a single moving resource. Future work could extend this scenario in several directions. First, exploring multi-resource scenarios could provide valuable insights into collective shepherding [65, 66] and collective hunting [34] behaviors. Second, investigating heterogeneous policies could allow for beneficial specialization among agents [67]. While our decentralized training experiments allowed for such heterogeneity, this primarily led to a decrease in coordination for the Cohesive Tracking strategy. A promising future direction would be to explore whether heterogeneity could instead lead to more efficient task distribution, such as explicit Producer and Scrounger roles, under different collective foraging scenarios. Third, our agents operated with significant informational constraints: they were memory-less (reacting only to the current time step) and perceived only one peer at a time, sampled uniformly from their visual field. Introducing memory, for instance, would allow agents to integrate information over time to learn resource trajectories or build reputations [68, 69]. Similarly, extending the model to incorporate distance-dependent attention mechanisms, allow for multi-sensor integration–such as combining visual and audio cues [70–72] (e.g., alarm or food recruitment calls) or aggregating information from multiple peers simultaneously [73]–would be a critical step toward understanding more complex attentional mechanisms and collective decision-making, better capturing the rich social dynamics observed in natural systems.

## Supporting information

ESM

## Data Accessibility

All the source code necessary to reproduce the simulation environment, multi-agent reinforcement learning training, and evaluation pipelines is available on Zenodo: https://doi.org/10.5281/zenodo.20817076. The accompanying dataset, which includes the trained model parameters and aggregated evaluation rollouts required to directly reproduce the figures and analysis without retraining, is available in a separate Zenodo repository: https://doi.org/10.5281/zenodo.20819723.

## Authors’ Contributions

V.C. and P.R. conceptualized the study. V.C. performed the simulations, curated the data, conducted the analysis, and prepared the figures under the supervision of P.R. V.C. wrote the original manuscript draft, and all authors contributed to reviewing and editing the final version.

## Acknowledgements

We are grateful to Dr. Mohsen Raoufi and Dr. Valentin Lecheval for fruitful discussions regarding the simulation design and for their insightful comments on the manuscript.

