## Supplementary material for "Social Information Quality and Environmental Volatility Shape Collective Foraging Behavior": ESM

### Contents of Supplementary Information

**S1 Sensitivity to visual range** **2**

**S2 Supplementary Figures** **3**

**S4 Supplementary Videos** **13**

### S1 Sensitivity to visual range

We performed additional analyses to assess how agent performance and learned policies are sensitive to variations in visual range ($r_{\text{vis}}$). The main results reported use a visual range of $r_{\text{vis}} = 15$. Fig. S1 shows how reducing the visual range to 10, 5, and 1 affects agent performance across all experimental conditions. Due to computational limitations, these results were not reproduced across multiple training seeds. The confidence intervals are calculated between policy evaluation steps and indicate how much the policy changes during the final phase of training. Performance results remain stable at visual ranges of 10 and 5, almost identically reproducing the results presented in the main text ( Fig. 2), and deteriorate only at the minimal range of 1, highlighting the robustness of our results to moderate changes in perceptual constraints. Robustness to moderate changes in visual range is expected, as the learned individual behavior for all conditions results in formation of compact groups. See Section 3.3 for more information.

An extremely short visual range makes training unstable, especially in the conditions when agents can benefit most from social information (Fig. S1, right part of the right panel). This instability likely stems from a violation of the synchronization guarantees established for blinking network models [38, 39]. In other words, agents observe others too infrequently or not randomly, so that the time aggregate no longer equals the instantaneous social information average. Alternatively, an $r_{vis} = 1$ could lead to extremely sparse social observations, preventing effective learning of the value of social cues.

### S2 Supplementary Figures

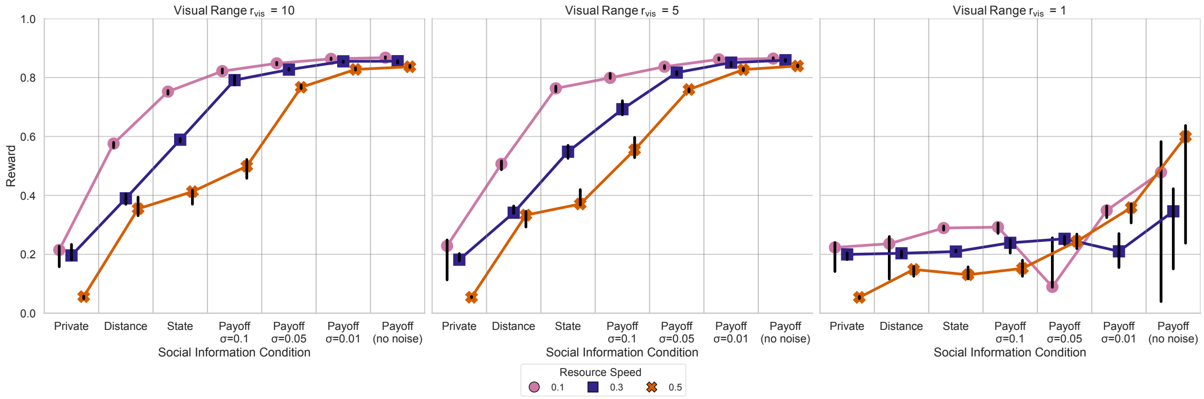

**Figure S1: Impact of visual range on performance across conditions.** Median normalized rewards (±95% CI) are shown for agents operating under seven social information conditions, across three resource speeds. Point markers indicate median reward. Social information quality conditions incrementally add features: starting from private information only, followed by access to (1) distance to a neighbor, (2) neighbor's current action (state), and (3) neighbor's reward signal with decreasing noise ($\sigma = 0.1$ , $\sigma = 0.05$, $\sigma = 0.01$ , and no noise). Marker shapes and bar colors indicate resource speeds (relative to the constant $v_{\max}$).

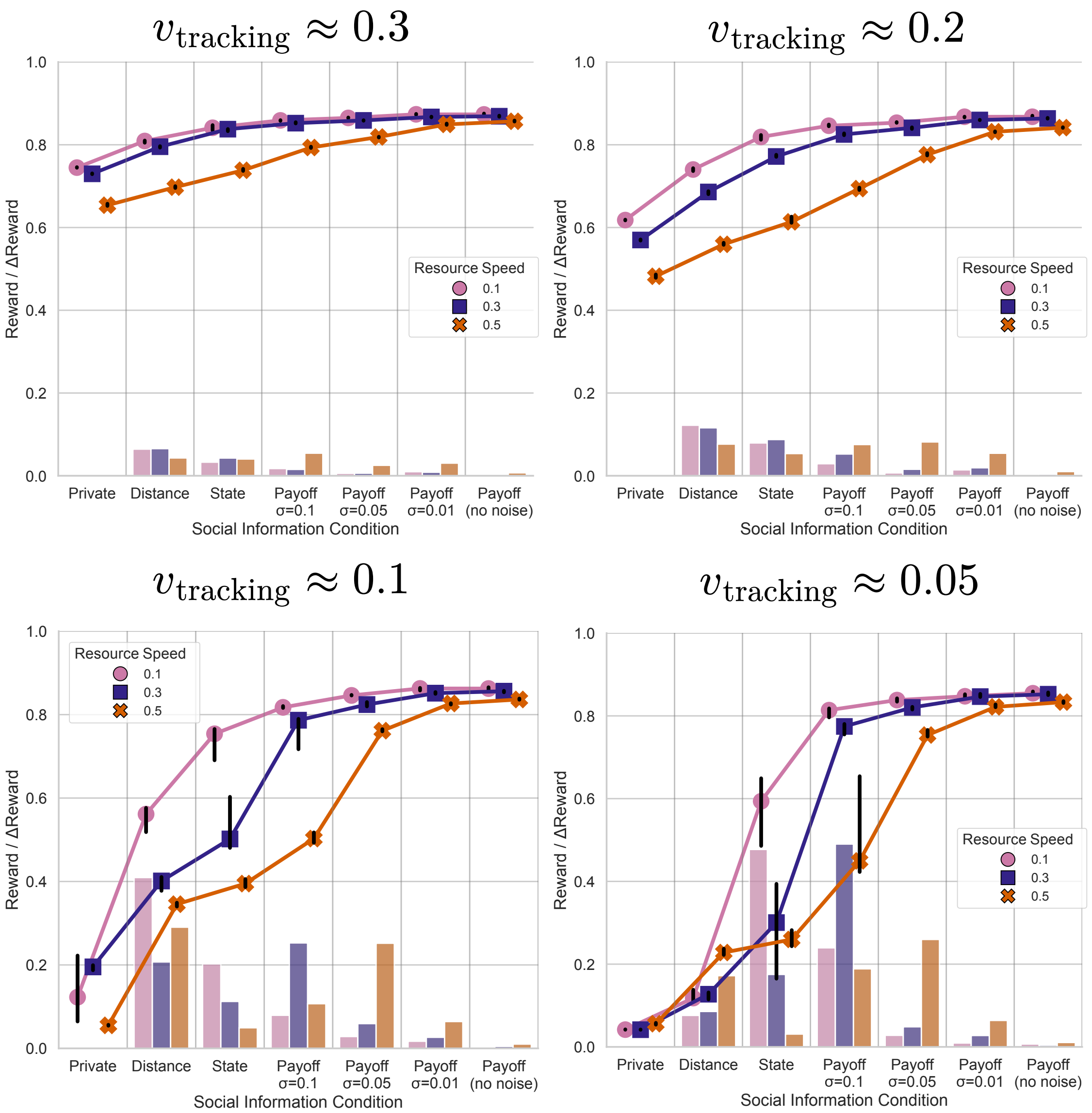

**Figure S2: Impact of effective tracking speed on performance.** Median normalized rewards (±95% CI) are shown as a function of social information quality and resource speed, analogous to Fig. 2 in the main text. Each panel corresponds to a different effective tracking speed ($v_{\text{tracking}}$). The speeds shown are $v_{\text{tracking}} = 0.3 \times v_{\max}$, $0.2 \times v_{\max}$, $0.1 \times v_{\max}$ (the value used in the main text), and $0.05 \times v_{\max}$ (corresponding to $p_{\text{track}}$ values of 0.3, 0.2, 0.1, and 0.05 respectively). Lower effective tracking speeds (i.e., higher intrinsic tracking costs) lead to a general decrease in performance, especially in low-quality social information conditions where agents rely heavily on the tracking action.

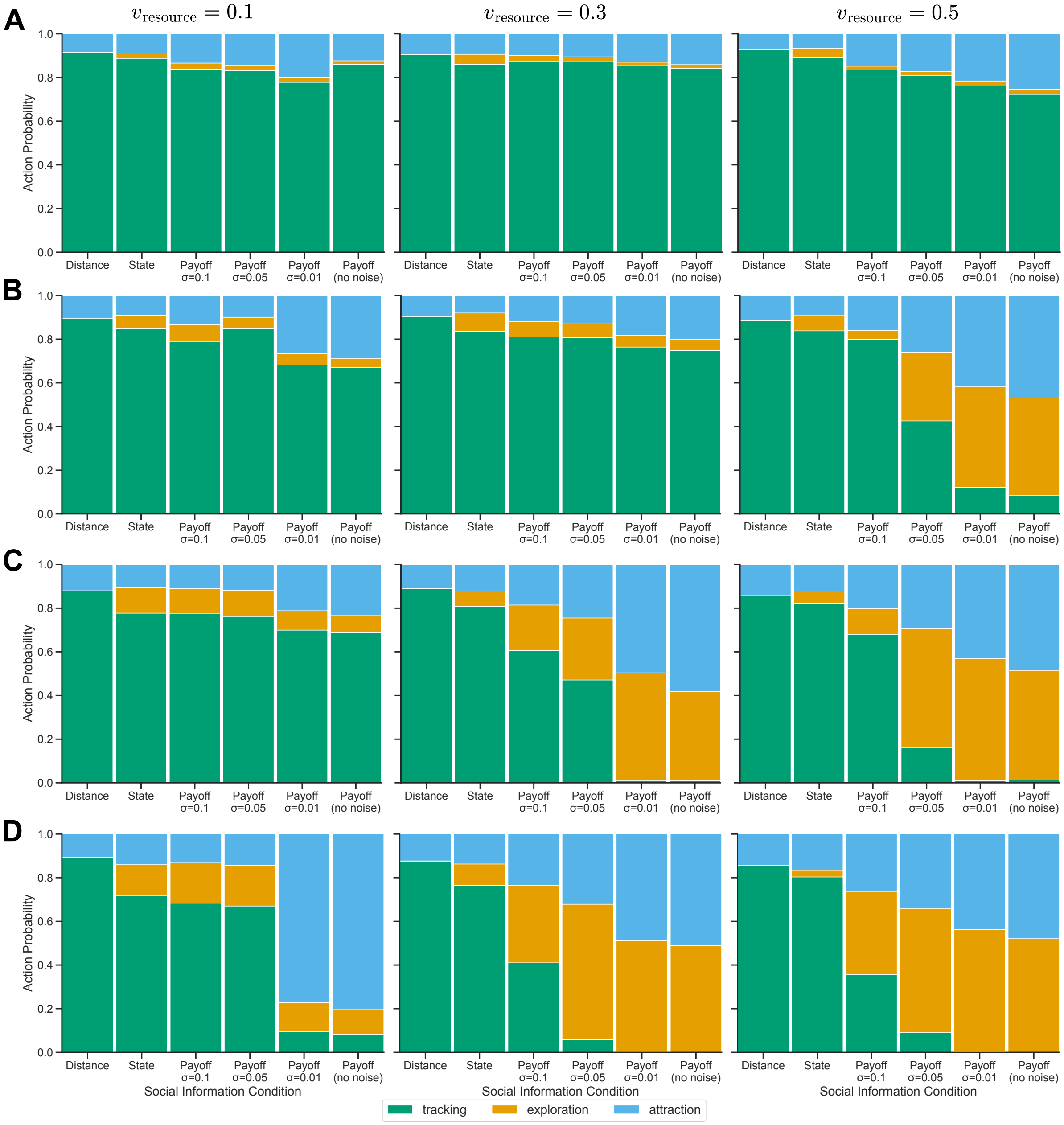

**Figure S3: Impact of effective tracking speed on behavioral strategy.** Average proportion of time agents spent in each behavioral mode (tracking, exploration, social attraction) across different effective tracking speeds. Each row corresponds to a different effective tracking speed ($v_{\text{tracking}}$): (**A**) $0.3 \times v_{\max}$, (**B**) $0.2 \times v_{\max}$, (**C**) $0.1 \times v_{\max}$ (the value used in the main text), and (**D**) $0.05 \times v_{\max}$. Within each row, panels correspond to different resource speeds (e.g., $v_{\text{resource}} = 0.1 \times v_{\max}$). Decreasing the effective tracking speed consistently reduces the reliance on tracking (green) and promotes exploration (orange) and social attraction (blue), especially in high-quality social information conditions.

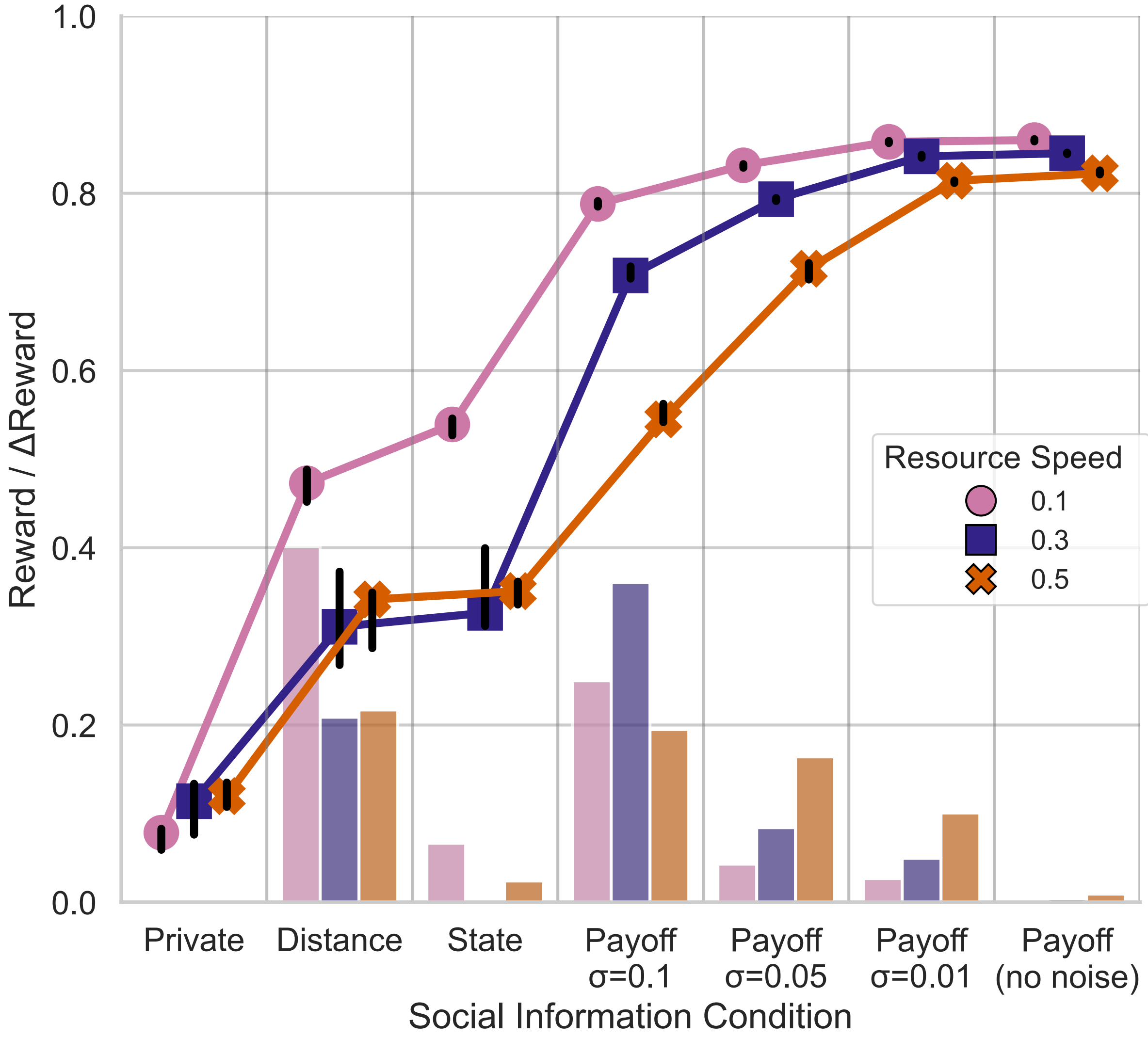

**Figure S4: Performance with decentralized training.** Median normalized rewards (±95% CI) are shown for agents trained with a fully decentralized version of MAPPO (i.e., without a centralized critic and actor networks, IPPO). The layout is analogous to Fig. 2 in the main text. Although the overall trends are similar to those of centralized training (Fig. 2), performance is notably lower under conditions of low-quality social information, which highlights the fragility of emergent strategies based on low-quality social cues.

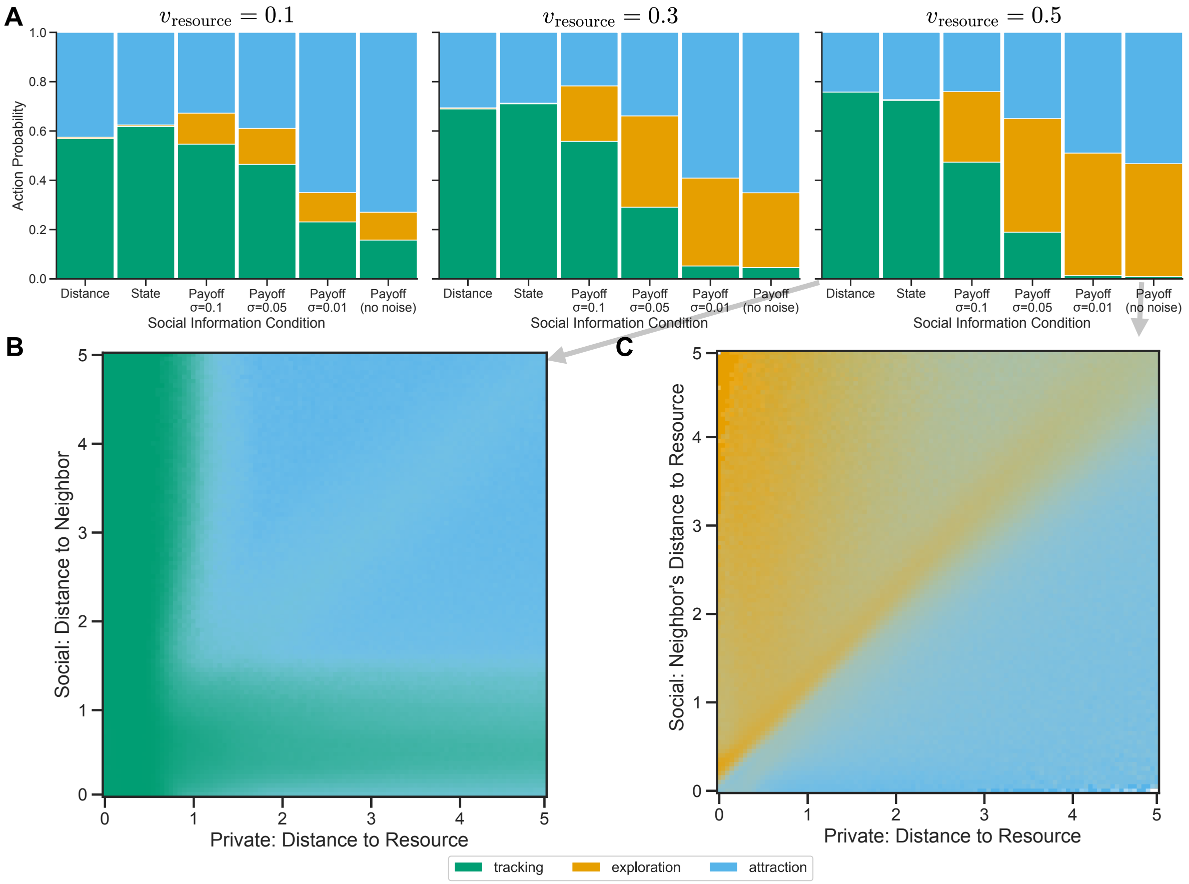

**Figure S5: Emergent strategies with decentralized training.** **A**: Average proportion of time agents spent in each behavioral mode across all conditions, analogous to Fig. 3. Compared to centralized training, agents show an increased reliance on social attraction. **B** and **C**: Emergent behavioral rules for the most volatile environment ($v_{\text{resource}} = 0.5 \times v_{\text{max}}$) under two extreme social information conditions, analogous to the left column of Fig. 4 **B**: In the "distance-only" social information condition, the Cohesive Tracking strategy is less pronounced, with a larger region of individual tracking. **C**: In the "noiseless payoff" social information condition, the Explore-or-Copy strategy remains dominant analogous to centralized training.

### S3 Supplementary Tables

**Table S1:** Third-party software assets

| **Package** | **Version** | **License** † | **License URL** |
|---|---|---|---|
| Python | 3.9 | PSF | docs.python.org |
| Hydra-Core | 1.3.2 | MIT | github.com/. . . /hydra |
| Matplotlib | 3.9.0 | PSF/BSD | matplotlib.org |
| Scikit-learn | 1.6.1 | BSD-3-Clause | github.com/.../scikit-learn |
| Seaborn | 0.13.2 | BSD-3-Clause | github.com/. . . /seaborn |
| wandb | 0.19.2 | MIT | github.com/. . . /wandb |
| PyTorch (cu121) | 2.5.1 | BSD-3-Clause | github.com/. . . /pytorch |
| TorchRL | 0.6.0 | MIT | github.com/. . . /rl |
| TensorDict | 0.6.2 | MIT | github.com/. . . /tensordict |
| VMAS | 1.4.3 | GPL-3.0 | github.com/. . . /VMAS |

Third-party software assets used in this work. All are installable via `pip` or `poetry`.
†Short identifiers follow the SPDX licence list.

**Table S2:** Training hyperparameters

| **Parameter** | **Value** |
|---|---|
| *Data Collection* | |
| Frames per batch | 500,000 |
| Number of iterations | 480 |
| Total training frames per condition | 240M |
| *Loss Function* | |
| Discount factor ($\gamma$) | 0.99 |
| GAE parameter ($\lambda$) | 0.9 |
| Entropy coefficient | 1e-4 |
| Clipping parameter ($\epsilon$) | 0.1 |
| *Optimization* | |
| Learning rate | $5 \times 10^{-5}$ |
| Number of epochs | 45 |
| Minibatch size | 4096 |
| Max gradient norm | 40.0 |
| *Network Architecture (Critic & Actor)* | |
| Hidden layers | 2 |
| Hidden units per layer | 256 |
| Activation function | Tanh |

Parameters were selected based on recommendations from the existing MARL literature, optimizing for stability and convergence of agent training across conditions [40–43]. In the supplementary code, these hyperparameters are listed in the `parameters.yaml` file along with the other settings for training and evaluation.

**Table S3:** Strategy consistency

| $v_{\text{resource}}$ | Social info | $N$ pruned seeds | All seeds | After pruning |
|---|---|---|---|---|
| 0.1 | Private | 2 | **0.778** [0.679, 0.866] | 0.934 [0.889, 0.971] |
| 0.1 | Distance | 0 | 1.000 [1.000, 1.000] | 1.000 [1.000, 1.000] |
| 0.1 | State | 0 | 0.990 [0.985, 0.994] | 0.990 [0.985, 0.994] |
| 0.1 | Payoff (no noise) | 1 | 0.913 [0.856, 0.963] | 0.989 [0.981, 0.995] |
| 0.1 | Payoff $\sigma$=0.01 | 0 | 0.966 [0.950, 0.980] | 0.966 [0.950, 0.980] |
| 0.1 | Payoff $\sigma$=0.05 | 0 | 0.999 [0.999, 0.999] | 0.999 [0.999, 0.999] |
| 0.1 | Payoff $\sigma$=0.1 | 0 | 0.999 [0.998, 0.999] | 0.999 [0.998, 0.999] |
| 0.3 | Private | 1 | **0.816** [0.689, 0.936] | 0.997 [0.995, 0.999] |
| 0.3 | Distance | 0 | 0.999 [0.998, 0.999] | 0.999 [0.998, 0.999] |
| 0.3 | State | 0 | 0.984 [0.978, 0.989] | 0.984 [0.978, 0.989] |
| 0.3 | Payoff (no noise) | 0 | 0.985 [0.977, 0.992] | 0.985 [0.977, 0.992] |
| 0.3 | Payoff $\sigma$=0.01 | 0 | 0.997 [0.996, 0.999] | 0.997 [0.996, 0.999] |
| 0.3 | Payoff $\sigma$=0.05 | 1 | 0.955 [0.928, 0.979] | 0.990 [0.985, 0.995] |
| 0.3 | Payoff $\sigma$=0.1 | 0 | 0.980 [0.969, 0.989] | 0.980 [0.969, 0.989] |
| 0.5 | Private | 0 | 0.991 [0.987, 0.994] | 0.991 [0.987, 0.994] |
| 0.5 | Distance | 0 | 0.998 [0.997, 0.999] | 0.998 [0.997, 0.999] |
| 0.5 | State | 0 | 0.990 [0.983, 0.995] | 0.990 [0.983, 0.995] |
| 0.5 | Payoff (no noise) | 0 | 0.990 [0.986, 0.994] | 0.990 [0.986, 0.994] |
| 0.5 | Payoff $\sigma$=0.01 | 0 | 0.998 [0.998, 0.999] | 0.998 [0.998, 0.999] |
| 0.5 | Payoff $\sigma$=0.05 | 0 | 0.997 [0.996, 0.998] | 0.997 [0.996, 0.998] |
| 0.5 | Payoff $\sigma$=0.1 | 0 | 0.979 [0.964, 0.992] | 0.979 [0.964, 0.992] |

Average pair-wise trajectory similarity (mean and bootstrap 95 % CI) for each social information condition and resource speed $v_{\text{resource}}$. Values are calculated over the upper triangle of the similarity matrices; the right-most column excludes training runs that were pruned. Average similarity below 0.9 are highlighted.

**Table S4:** Rewards

| $v_{\mathrm{resource}}$ | Social info | Reward | Δ Reward |
|---|---|---|---|
| 0.1 | Private | 0.123 [0.065, 0.222] | – |
| 0.1 | Distance | 0.561 [0.518, 0.576] | **0.409** [0.367, 0.424] |
| 0.1 | State | 0.754 [0.691, 0.765] | **0.202** [0.139, 0.214] |
| 0.1 | Payoff $\sigma$=0.1 | 0.817 [0.816, 0.820] | 0.079 [0.078, 0.082] |
| 0.1 | Payoff $\sigma$=0.05 | 0.846 [0.846, 0.847] | 0.028 [0.028, 0.030] |
| 0.1 | Payoff $\sigma$=0.01 | 0.863 [0.858, 0.864] | 0.017 [0.012, 0.018] |
| 0.1 | Payoff (no noise) | 0.862 [0.858, 0.866] | 0.001 [-0.004, 0.004] |
| 0.3 | Private | 0.195 [0.187, 0.199] | – |
| 0.3 | Distance | 0.401 [0.378, 0.411] | **0.207** [0.184, 0.217] |
| 0.3 | State | 0.502 [0.481, 0.603] | 0.113 [0.091, 0.213] |
| 0.3 | Payoff $\sigma$=0.1 | 0.787 [0.717, 0.790] | **0.253** [0.183, 0.256] |
| 0.3 | Payoff $\sigma$=0.05 | 0.824 [0.820, 0.830] | 0.059 [0.055, 0.065] |
| 0.3 | Payoff $\sigma$=0.01 | 0.851 [0.850, 0.854] | 0.027 [0.025, 0.029] |
| 0.3 | Payoff (no noise) | 0.856 [0.854, 0.857] | 0.005 [0.002, 0.006] |
| 0.5 | Private | 0.055 [0.054, 0.056] | – |
| 0.5 | Distance | 0.346 [0.340, 0.349] | **0.291** [0.285, 0.294] |
| 0.5 | State | 0.395 [0.387, 0.406] | 0.049 [0.041, 0.060] |
| 0.5 | Payoff $\sigma$=0.1 | 0.503 [0.493, 0.518] | 0.107 [0.097, 0.122] |
| 0.5 | Payoff $\sigma$=0.05 | 0.762 [0.760, 0.765] | **0.252** [0.250, 0.256] |
| 0.5 | Payoff $\sigma$=0.01 | 0.827 [0.824, 0.827] | 0.064 [0.062, 0.065] |
| 0.5 | Payoff (no noise) | 0.836 [0.836, 0.838] | 0.010 [0.010, 0.012] |

Median normalized reward and reward change (Δ Reward) with 95 % bootstrap confidence intervals. Δ Rewards above 0.2 are highlighted.

**Table S5:** States

| $v_{\text{resource}}$ | Social Info | Attraction | Tracking | Exploration |
|---|---|---|---|---|
| 0.1 | Distance | 0.120 [0.118, 0.123] | **0.879** [0.877, 0.882] | 0.000 [0.000, 0.000] |
| 0.1 | State | 0.107 [0.094, 0.121] | **0.777** [0.739, 0.818] | 0.116 [0.063, 0.165] |
| 0.1 | Payoff $\sigma$=0.1 | 0.110 [0.107, 0.114] | **0.774** [0.760, 0.788] | 0.116 [0.099, 0.132] |
| 0.1 | Payoff $\sigma$=0.05 | 0.118 [0.109, 0.128] | **0.762** [0.750, 0.776] | 0.120 [0.110, 0.131] |
| 0.1 | Payoff $\sigma$=0.01 | 0.212 [0.143, 0.287] | **0.700** [0.624, 0.769] | 0.089 [0.086, 0.090] |
| 0.1 | Payoff (no noise) | 0.234 [0.192, 0.282] | **0.688** [0.640, 0.731] | 0.078 [0.073, 0.082] |
| 0.3 | Distance | 0.110 [0.086, 0.127] | **0.890** [0.873, 0.914] | 0.000 [0.000, 0.000] |
| 0.3 | State | 0.121 [0.099, 0.140] | **0.807** [0.748, 0.868] | 0.071 [0.020, 0.133] |
| 0.3 | Payoff $\sigma$=0.1 | 0.186 [0.180, 0.191] | **0.606** [0.555, 0.658] | 0.209 [0.160, 0.256] |
| 0.3 | Payoff $\sigma$=0.05 | 0.245 [0.231, 0.257] | **0.471** [0.441, 0.504] | 0.284 [0.251, 0.307] |
| 0.3 | Payoff $\sigma$=0.01 | **0.496** [0.481, 0.514] | 0.011 [0.006, 0.018] | **0.493** [0.474, 0.511] |
| 0.3 | Payoff (no noise) | **0.581** [0.537, 0.625] | 0.010 [0.006, 0.013] | **0.409** [0.368, 0.451] |
| 0.5 | Distance | 0.141 [0.115, 0.157] | **0.859** [0.843, 0.885] | 0.000 [0.000, 0.000] |
| 0.5 | State | 0.122 [0.109, 0.133] | **0.823** [0.770, 0.864] | 0.055 [0.019, 0.108] |
| 0.5 | Payoff $\sigma$=0.1 | 0.202 [0.195, 0.208] | **0.681** [0.615, 0.725] | 0.118 [0.078, 0.178] |
| 0.5 | Payoff $\sigma$=0.05 | 0.295 [0.292, 0.297] | 0.160 [0.142, 0.177] | **0.546** [0.528, 0.565] |
| 0.5 | Payoff $\sigma$=0.01 | **0.430** [0.417, 0.442] | 0.010 [0.006, 0.014] | **0.560** [0.547, 0.575] |
| 0.5 | Payoff (no noise) | **0.485** [0.449, 0.518] | 0.013 [0.010, 0.016] | **0.503** [0.470, 0.536] |

Proportion of time agents spent in **Social Attraction**, **Tracking**, and **Exploration** states. Means with 95 % bootstrap confidence intervals for each state is presented. Proportions above 0.4 are highlighted.

### S4 Supplementary Videos

**Video 1: Cohesive Tracking.** This video shows the emergent strategy observed in a volatile environment ($v_{\text{resource}} = 0.5 \times v_{\text{max}}$) when agents have access only to low-quality social information (neighbor distance only). Agents are predominantly relying on private tracking while using social attraction non-selectively to maintain a compact, cohesive group cluster. Colors indicate the agents' current behavioral states: tracking (green), exploration (orange), or social attraction (blue).

**Video 2: Track-or-Copy.** This video shows the emergent strategy observed with high-quality payoff information when private tracking remains a viable default behavior (e.g., in conditions with a high effective tracking speed, $v_{\text{tracking}} = 0.3 \times v_{\text{max}}$, within a volatile resource environment, $v_{\text{resource}} = 0.5 \times v_{\text{max}}$). Agents default to private tracking and use social information selectively to copy peers with higher observed success. Colors indicate the agents' current behavioral states: tracking (green), exploration (orange), or social attraction (blue).

**Video 3: Explore-or-Copy.** This video shows the emergent strategy observed with high-quality payoff information in a volatile environment ($v_{\text{resource}} = 0.5 \times v_{\text{max}}$) where private tracking is rendered ineffective due to low effective tracking speed, $v_{\text{tracking}} = 0.05 \times v_{\text{max}}$. Agents abandon costly private tracking, default to random exploration, and opportunistically copy more successful neighbors. Colors indicate the agents' current behavioral states: tracking (green), exploration (orange), or social attraction (blue).
